# CXCR4 controls movement and degranulation of CD8^+^ T cells in the influenza-infected lung via differential effects on interaction and tissue scanning

**DOI:** 10.1101/2022.09.13.507813

**Authors:** Paulus Mrass, Ichiko Kinjyo, Janie R. Byrum, David Torres, Steven F. Baker, Judy L. Cannon

**Affiliations:** Department of Molecular Genetics and Microbiology, University of New Mexico School of Medicine, MSC 08 4660, 1 University of New Mexico, Albuquerque, NM 87131, USA; University of New Mexico Comprehensive Cancer Center, University of New Mexico Health Sciences Center, Albuquerque, NM, USA; Department of Internal Medicine, Division of Molecular Medicine, University of New Mexico Health Sciences Center, Albuquerque, NM, USA; Department of Math and Physical Science, Northern New Mexico College, Espanola, NM; Infectious disease Program, Lovelace Biomedical Research Institute, Albuquerque, NM; Autophagy, Inflammation, and Metabolism Center of Biomedical Research Excellence, University of New Mexico School of Medicine, Albuquerque NM

## Abstract

Effector CD8^+^ T cell interactions are critical in controlling viral infection by directly killing infected cells but overabundant or sustained activation also exacerbates tissue damage. Chemokines promote the trafficking of effector CD8^+^ T cells into infected tissues, but we know little about how chemokines regulate the function of CD8^+^ T cells within tissues. Using a murine model of influenza A virus infection, we found that expression of the chemokine receptor CXCR4 by lung-infiltrating cytotoxic T cells correlated with the expression of the degranulation marker CD107a. Inhibition of CXCR4 reduced activation, adhesion, and degranulation of cytotoxic T cells *in vitro* and *in vivo*. Moreover, in live influenza-infected lung tissue, T cells stopped moving in lung regions with high levels of influenza antigen, and CXCR4 was essential for CD8^+^ T cells to execute this arrest signal fully. In contrast, CXCR4 increased the motility of CD8^+^ T cells in low-influenza areas of the lung. We also found that CXCR4 stimulated the effector function of lung-infiltrating cytotoxic T cells even after clearance of influenza virus, and inhibition of CXCR4 expedited the recovery of influenza-infected mice, despite delayed clearance of the replication-competent virus. Our results suggest that CXCR4 promotes the interaction strength of cytotoxic T cells in lung tissue through combined effects on T cell movement and interaction with virally infected target cells in influenza infected-lungs.

## Introduction

CD8^+^ effector T cells are essential for effective clearance of viral infections, including influenza A virus (Cullen et al., 2019; Duan and Thomas, 2016; Hufford et al., 2015). Yet prolonged activation and persistence of CD8^+^ T cells can also cause damage to peripheral tissue after the elimination of the infection (Duan and Thomas, 2016; Goplen et al., 2020; Hufford et al., 2015). Effective T cell function requires migration and interaction with target cells in the lymph node and peripheral tissues. In response to infection, interactions between naïve CD8^+^ T cells and dendritic cells in the lymph nodes induce priming and differentiation into virus-specific effector cytotoxic CD8^+^ T cells. These differentiated effector T cells then migrate into infected tissue, where they directly interact with virally infected cells. Interactions with infected target cells lead to the release of cytotoxic granules and consequently the killing of virally infected cells, which contributes to the elimination of the infection.

Interactions between T cells and target cells regulate T cell differentiation and effector function. Recent studies have shown that fine-tuning the contact durations between T cells and dendritic cells in the lymph node impacts the differentiation pathway of CD4^+^ T cells (Groom et al., 2012; Mempel et al., 2004). Cytotoxic T cells infiltrating inflammatory tissue typically show sustained, stable contact with target cells (Deguine et al., 2010). Nevertheless, quantitative live imaging of cytotoxic T cells in virally infected skin suggested that cumulative short contacts by distinct cytotoxic T cells can also lead to the killing of target cells (Halle et al., 2016). However, we have an incomplete understanding of the molecules that regulate interaction duration and how fine-tuning contact durations might impact on the biological activity of cytotoxic T cells in infected tissues.

Chemokines are critical regulators of T cell migration and interaction at various stages of T cell function, from development to activation, differentiation, and effector function. For example, CCR7 is crucial for inducing naïve T cell movement in lymph nodes (Asperti-Boursin et al., 2007; Forster et al., 2008; Fricke et al., 2016; Letendre et al., 2015; Worbs and Forster, 2007), and CXCR3 and CCR5 promote contact durations between naïve T cells and dendritic cells within the lymph node, contributing to T cell differentiation (Castellino et al., 2006; Groom et al., 2012). In addition, several chemokine receptors induce T cell migration into infected tissues. For example, CXCL10 promotes CD4^+^ T cells to migrate into perivascular niches within the inflamed skin (Bala et al., 2022; Fowell and Kim, 2021; Gaylo-Moynihan et al., 2019; Prizant et al., 2021). Moreover, chemokines regulate T cell function during infections of the lung. For example, the influenza-infected lung upregulates the expression of the CXCR3 ligand CXCL10 (Ichikawa et al., 2013), which expedites the recruitment of lymphocytes into the infected lung tissue (Wareing et al., 2004). In addition, within the influenza-infected trachea, neutrophils generate CXCL12 trails that bind to CXCR4 on cytotoxic T cells, which promotes migration towards the epithelium of the influenza-infected trachea (Lim et al., 2015). In contrast, we know less about whether chemokine receptors directly regulate effector T cell function within influenza-infected lungs.

In this paper, we studied CXCR4 in a murine model of influenza to determine its role in primary immune responses against influenza. Our findings identify CXCR4 as a positive regulator of effector CD8^+^ T cell activation within influenza-infected lungs. Our data show that CXCR4 promotes degranulation of effector CD8^+^ T cells and enhances the duration of cytotoxic T cell interactions within the infected lung. Furthermore, CXCR4 promotes CD8^+^ effector function even after viral clearance, which may delay the recovery of influenza-infected mice. Our results support a role for CXCR4 in balancing cytotoxic T cells’ protective and immunopathogenic effects during respiratory tract infections.

## Results

### Lung-infiltrating CD8^+^ T cells express CXCR4 during anti-influenza immune responses

We used a murine model of influenza A virus (IAV) infection to determine the mechanisms that regulate cytotoxic T cell activation within the lungs (**Figure 1A**). We infected mice with the influenza A X31 strain, which causes transient disease as measured by weight loss, followed by rapid recovery (**Figure 1B).** Our primary focus was extravascular effector CD8^+^ T cells within the influenza-infected lung parenchyma. Therefore, we labeled intravascular T cells by intravenous injection of a CD3-specific antibody. This way, we could distinguish extravascular and intravascular T cells (Anderson et al., 2012). We found that effector CD8^+^ T cells accumulated in high numbers in the infected lung (**Figure 1C**). After day four post-infection, most CD8^+^ T cells were extravascular in the interstitial lung tissue (**Figure 1D**). To study the role of chemokine receptors in effector CD8^+^ T cell function, we measured expression levels of CXCR3 and CXCR4 starting at day six, when T cells migrate in significant numbers into the lung (**Figure 1D**; (Keating et al., 2018)). Previous reports supported CXCR3 and CXCR4 as key chemokine receptors that promote T cell recruitment into the lung and control T cell positioning within the trachea (Dhume et al., 2019; Hufford et al., 2015; Lim et al., 2015). We found that CXCR3 expression was close to background levels on day six after infection and increased only later in infection, becoming maximal on day ten, when mice already have recovered as measured by weight gain (**Figure 1E** and **1G**). In contrast, intravascular and extravascular cytotoxic T cells expressed CXCR4 consistently throughout the infection (**Figure 1F-H**). This expression pattern suggested that CXCR4 may play a role in CD8^+^ T cell function during the effector phase of an influenza-specific cytotoxic T cell response.

**Figure 1.**
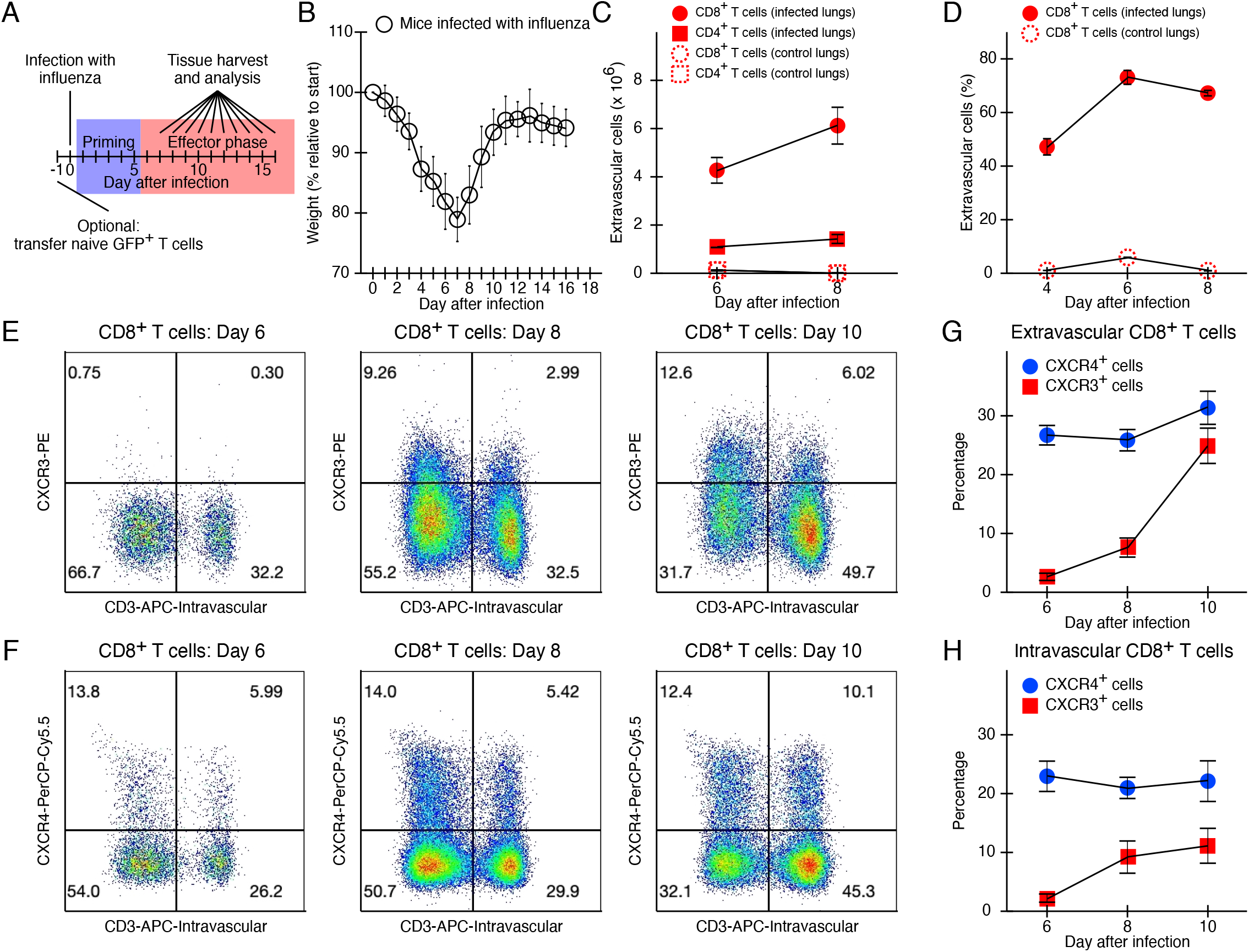
CXCR4 is stably expressed by cytotoxic T cells infiltrating influenza-infected lungs. (A) Experimental design. (B - H) C57BL/6 mice were infected with 1 x 10^3^ EID50 X31 influenza, followed by weight measurements (B) or flow cytometry analysis of lung tissue (C-H). (B) Weight measurements from two pooled independent experiments with a total of ten mice. The symbols indicate mean weight. The error bars show standard deviations. (C, D) Lung tissues from influenza-infected mice or uninfected control mice were analyzed with flow cytometry for number (C) and percentage (D) of extravascular T cells as determined by intravascular injection of anti-CD3 antibody five minutes before sacrifice. Shown is a representative experiment with two mice at each timepoint. Symbols indicate means and errors indicate standard errors. (E-H) The percentages of CXCR4^+^ and CXCR3^+^ cells were determined for intravascular and extravascular CD8^+^ T cells. Representative images are shown for CXCR3 (E) and CXCR4 (F) stains. Quantification of CXCR3^+^ and CXCR4^+^ T cells of extravascular (G) and intravascular (H) CD8^+^ T cells. Means and standard errors are shown (n = 6-7 mice at each time point, pooled from 4 independent experiments).

### CXCR4 expression correlates with degranulation of lung-infiltrating T cells

We next used the X31-OVA influenza model to analyze an antigen-specific OT-I T cell population that recognizes the OVA-influenza-associated antigen with high affinity (Thomas et al., 2006). We transferred naive GFP^+^ TCR-transgenic OT-I CD8^+^ T cells into recipient mice, followed by infection with the cognate X31-OVA strain. Infection with X31-OVA caused transient weight loss, similar to the native X31 strain (**data not shown**). Flow cytometry revealed that most effector OT-I CD8^+^ T cells in the lung were in the extravascular space, similar to infection with native X31 strain (**Figure 2A**). Approximately one-third of the effector OT-I T cells expressed CXCR4 (**Figure 2B**).

**Figure 2.**
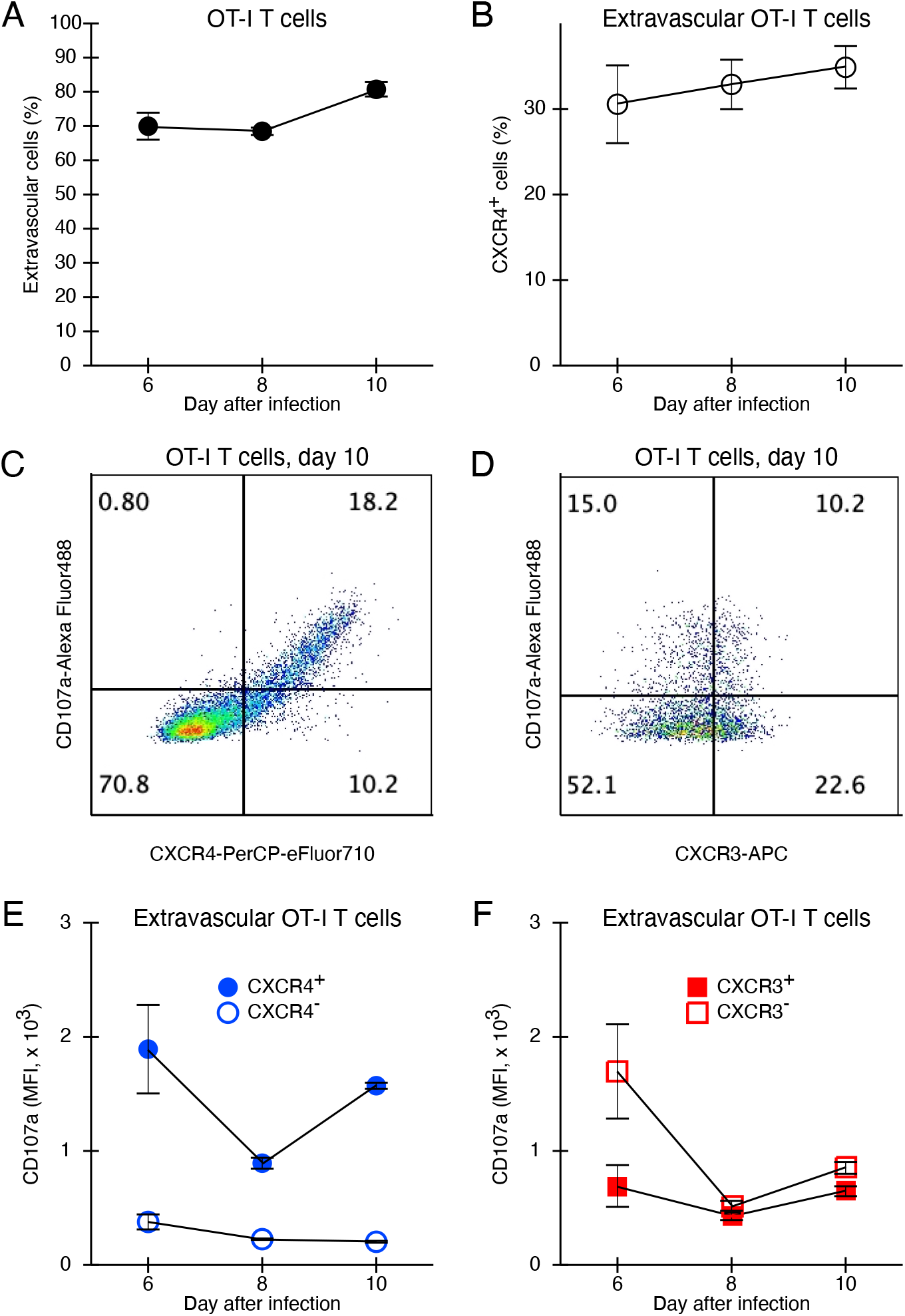
CXCR4 and CD107a surface expression are strongly correlated in cytotoxic T cells infiltrating influenza-infected lungs. (A - F) C57BL/6 mice received 1 x 10^4^ naive OT-I T cells, followed by infection with 1 x 10^5^ PFU X31-OVA influenza one day later. On day six, eight and 10 post-infection, single-cell suspensions were generated from lungs obtained from infected mice. The cell suspensions were stained to detect OT-I T cells and surface levels of CD107a and CXCR4. (A, B) The majority of OT-I T cells have extravasated (A) and approximately a third of extravascular T cells express CXCR4 on the cell surface (B). (C, D) Representative images for CXCR4/CD107a (C) and CXCR3/CD107a (D) stains. (E, F) Quantification of mean fluorescence intensity of CD107a depending on CXCR4 expression status (E) or CXCR3 expression status (F). Shown are means and standard errors from one representative experiment (n = 2, 3 mice per time point).

To obtain insight into a possible role of CXCR4 in regulating T cell effector functions within the lung environment, we stained lung-infiltrating T cells with an antibody for CD107a, a marker of recent degranulation (Betts et al., 2003). We found that CXCR4 expression correlated strongly with cytotoxic granule release (**Figure 2C**); effector CD8^+^ T cells high in CXCR4 show high levels of CD107a, while CXCR4-negative T cells have little CD107a (**Figure 2E**). In contrast, CXCR3 and CD107a showed no correlation, with CXCR3^+^ and CXCR3^-^ T cells showing similar levels of CD107a (**Figure 2D** and **2F**). These results suggest a correlation between CXCR4 expression and CD8^+^ effector T cell function.

### CXCR4 promotes the effector function of cytotoxic T cells

Previous studies have shown that in Jurkat cells, TCR triggering leads to a transfer of CXCR4 from the cell surface to the intracellular aspect of the immunological synapse, where it enhances TCR-dependent signaling (Kumar et al., 2006). To test whether CXCR4 has a similar role in primary cytotoxic T cells, we analyzed CXCR4-deficient cytotoxic T cells in well-defined *in vitro* assays. We generated CXCR4-deficient cytotoxic T cells by *in vitro* expansion of CD8^+^ T cells obtained from CD4-Cre : CXCR4^Flox/Flox^ mice (CXCR4^KO^). Cytotoxic T cells from CXCR4^Flox/Flox^ littermates that did not express CD4-Cre (CXCR4^WT^) served as controls. We found that TCR-crosslinking downregulates CXCR4 from the surface of wild-type CXCR4^WT^ cytotoxic T cells (**Figure 3A**). Moreover, CXCR4-deficient T cells showed decreased calcium mobilization in response to TCR-crosslinking (**Figure 3B**). These results demonstrate that CXCR4 promotes TCR-signaling in primary cytotoxic T cells.

**Figure 3.**
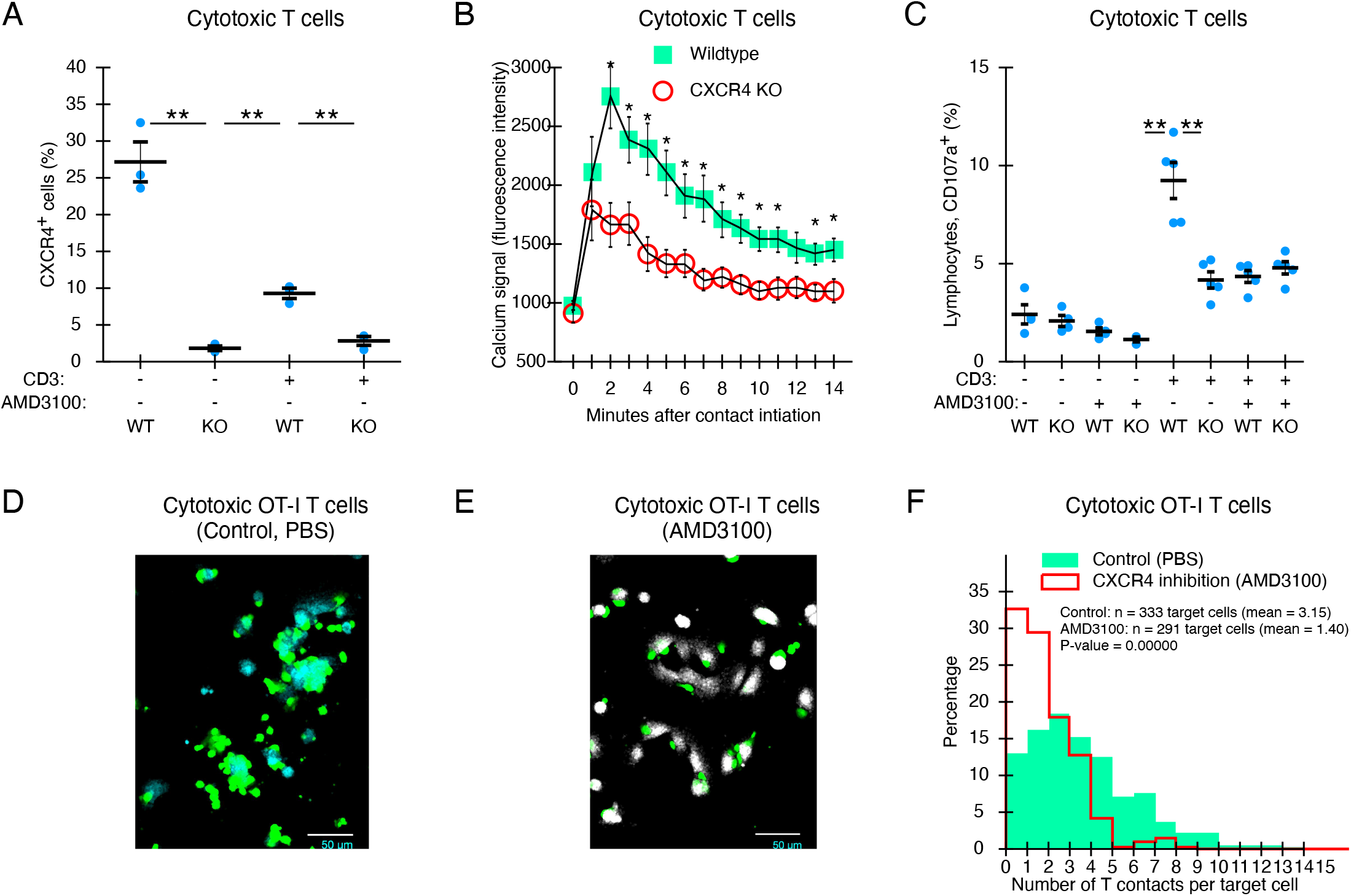
CXCR4 promotes TCR-dependent activation of cytotoxic T cells *in vitro*. (A – C) We isolated naïve CD8^+^ T cells from spleens from CD4-Cre : CXCR4^Flox/Flox^ (CXCR4^KO^) mice and littermates that did not express the CD4-Cre transgene (CXCR4^WT^). We stimulated CD8^+^ T cells with plate-bound anti-CD3 and anti-CD28 antibodies, and expanded them with rhIL-2. Between four and seven days after stimulation, T cells were re-seeded on flat-bottom 96 wells coated with anti-CD3 antibody. (A, C) After 2 hours of incubation, we stained T cells with an antibody for CXCR4 (A) or CD107a (C), measured fluorescence intensity with the BD Fortessa Flow Cytometer, and quantified the percentage of CXCR4^+^ and CD107a^+^ T cells with FlowJo. The graphs represent individual experiments, and the symbols represent single wells. (B) T cells were loaded with the calcium-sensitive fluorophore Cal-520, then imaged using the Incucyte Live Cell Imager with a frame rate of one image per minute. The mean fluorescence of individual T cells was measured from the time when they made contact with the bottom of the dish coated with anti-CD3 antibody. Each symbol represents the mean value +/- SEM of 15 T cells. Experiments were repeated 3 times. P-values were calculated with the Student’s t-test (*: <0.05; **: <0.005.) (D – F) CD8^+^ T cells from splenocytes of B6Ub-GFP x OT-I mice were isolated, stimulated with OVA and seeded on culture dishes with plated ID8-RFP-OVA cells. To inhibit CXCR4, we added 50 μM AMD3100, and fixed the cells after overnight incubation. Cells were imaged with confocal microscopy after washing. We quantified the number of T cells that contacted each ID8-RFP-OVA target cells and the graph shows the number of T cell contacting each ID8-OVA target cell. Data pooled from three independent experiments. The p-value was calculated with the Mann-Whitney test.

We then determined the role of CXCR4 in TCR crosslinking-induced degranulation of cytotoxic T cells by measuring the surface expression of CD107a in CXCR4^KO^ CD8^+^ T cells. We found that CXCR4^KO^ CD8^+^ T cells showed significantly less degranulation than CXCR4^WT^ T cells (**Figure 3C**). We then used a well-established inhibitor of CXCR4, AMD3100, to confirm the role of CXCR4 on degranulation. AMD3100 is used clinically as a specific CXCR4 antagonist (Wang et al., 2020). We found that CXCR4^WT^ CD8^+^ T cells treated with the CXCR4 inhibitor AMD3100 had similar levels of CD107a as CXCR4^KO^ CD8^+^ T cells (**Figure 3C**). In addition, AMD3100 did not further affect the already reduced degranulation of CXCR4^KO^ T cells (**Figure 3C**). These findings demonstrated that AMD3100 is a specific inhibitor of CXCR4 in cytotoxic T cells and that CXCR4 directly stimulates CD8^+^ T cell degranulation.

Chemokine receptors, including CXCR4, regulate T cell movement and adhesion to the endothelium (Alon et al., 2021; Scimone et al., 2004). To test whether CXCR4 can directly affect T cell adhesion, we quantified the number of contacts between OVA-specific cytotoxic OT-I T cells and target cells that express OVA, ID8-OVA with and without CXCR4 inhibitor AMD3100. We found that treatment with the CXCR4 inhibitor AMD3100 reduced the number of T cells interacting with plated ID8-OVA cells (**Figure 3D-F**). This result shows that CXCR4 enhances TCR-induced T cell adhesiveness. Together, these data indicate that CXCR4 enhances cognate TCR-signaling in cytotoxic CD8^+^ T cells and downstream effects, including degranulation and adhesion.

### CXCR4 increases degranulation of cytotoxic T cells *in vivo* in influenza-infected lungs

Next, we asked whether CXCR4 regulates the effector function of cytotoxic T cells in infected lungs. We transferred naïve CD8^+^ OT-I T cells into recipient mice and infected the animals with X31-OVA. To minimize potential effects of CXCR4 on T cell priming and recruitment into the lung, we injected animals with 200 μg of AMD3100 via intraperitoneal injection every twelve hours starting from day five post-infection, i.e., the timepoint when T cells begin to accumulate in the lung (Keating et al., 2018) (**Figure 1**). This treatment regimen decouples the effects on CXCR4 inhibition on T cell activation and recruitment to the lung from CXCR4 effects on effector T cells in the lung after infection. Flow cytometry analysis of lungs on days seven and ten after infection showed that the absolute numbers of OT-I T cells were similar in AMD3100-treated and untreated animals (**Figures 4A and 4E**). Similarly, T cells extravasated effectively after inhibition of CXCR4 starting on day five (**Figures 4B and 4F**).

**Figure 4.**
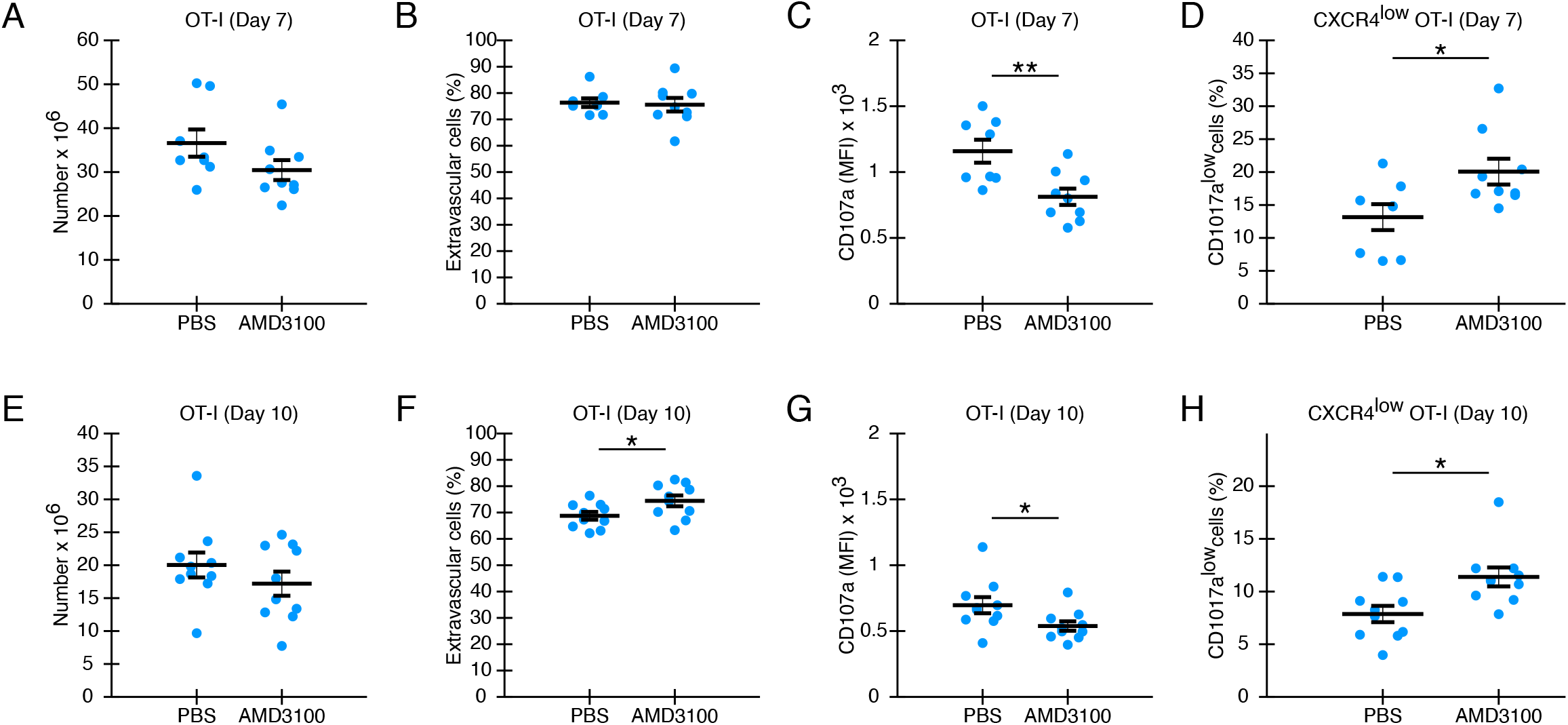
Inhibition of CXCR4 reduces cytotoxic T cell degranulation in influenza-infected lungs. C57BL/6 mice received 1 x 10^4^ naive GFP+ OT-I T cells, followed by infection with 1 x 10^5^ PFU X31-OVA influenza. On day 7 (A-D) and day 10 (E-H) post-infection, single-cell suspensions of the lungs were generated and analyzed with flow cytometry. Before tissue harvest, mice received a fluorescently labeled CD3 antibody through the tail-vein to label intravascular T cells. (A and E) show the total numbers of GFP+ OT-I T cells in the lungs. (B and F) show the percentage of OT-I T cells that have extravasated into lung tissue. (C and G) show the mean CD107a fluorescence of extravasated OT-I T cells. (D and H) show the percentage CD107a^low^ T cells of extravasated OT-I T cells that express low-medium levels of CXCR4. Each symbol shows the values obtained from an individual mouse. Data from two independent experiments were pooled (day 7: n = 8 mice in the control group; 9 mice in the AMD3100 group; day 10: n = 10 mice in either group). Lines indicate the mean and errors show standard error of the mean. P-values were calculated with the Student’s t-test. (*: <0.05; **:<: 0.005).

We then analyzed CD8^+^ effector function after CXCR4 inhibition in the influenza-infected lung by quantifying the fluorescence intensity of CD107a. We found reduced CD107a expression on CD8^+^ T cells after inhibition of CXCR4 on day seven post-infection (**Figure 4C**). Furthermore, reduced CD107a expression persisted until day ten (**Figure 4G**), after the clearance of the virus. Because we found a correlation between CXCR4 expression and CD107a, we also quantified CD107a on OT-I T cells in T cells with low expression levels of CXCR4 (CXCR4^low^). We found that in the CXCR4^low^ population, CXCR4 inhibition led to an increased percentage of CD107a low cells on both days seven (**Figure 4D**) and ten (**Figure 4H**) post-infection. Thus, even after normalizing the CXCR4 expression level, CXCR4-inhibition in mice decreased CD107a on the surface of lung-infiltrating T cells. Together, these data show that CXCR4 promotes degranulation by cytotoxic T cells in influenza-infected lungs at the peak of infection and after viral clearance.

### Cytotoxic T cells in the lung switch between confinement and ballistic relocation

Because CXCR4 strengthened TCR-induced adhesions *in vitro* (**Figure 3**), we asked whether CXCR4 played a similar role in the influenza-infected lung tissue. As a readout for adhesiveness in virally infected lung tissue, we quantified the duration of migration stops using a previously established live tissue imaging model of inflamed lungs (Mrass et al., 2017). We first generated a detailed measurement of cytotoxic T cell migration in influenza-infected lungs. Briefly, we adoptively transferred naïve polyclonal GFP^+^ CD8^+^ T cells into recipient mice, followed by infection with X31. We removed influenza-infected lungs at days six to eight post-infection and imaged GFP^+^CD8^+^ effector T cells responding to influenza infection within live lung tissue.

Careful inspection of the movement of CD8^+^ T cells in influenza-infected lungs revealed transitions between active migration and confinement (**Movie 1**). This pattern was similar to the previously described intermittent migration pattern for non-antigen-specific effector CD8^+^ T cells in LPS-inflamed lung tissue (Mrass et al., 2017). To quantify active movement and stop phases of effector CD8^+^ T cells in influenza-infected lungs, we measured how long individual T cells stayed consistently within a 15 μm radius from their origin and called this parameter the “dwell time”. **Figure 5A** shows migration phases with short dwell times highlighted in green (“go” phase) and phases with long dwell times highlighted in red (“stop” phase). This depiction indicates that active translocation in space happens almost exclusively when T cells show short dwell times.

**Figure 5.**
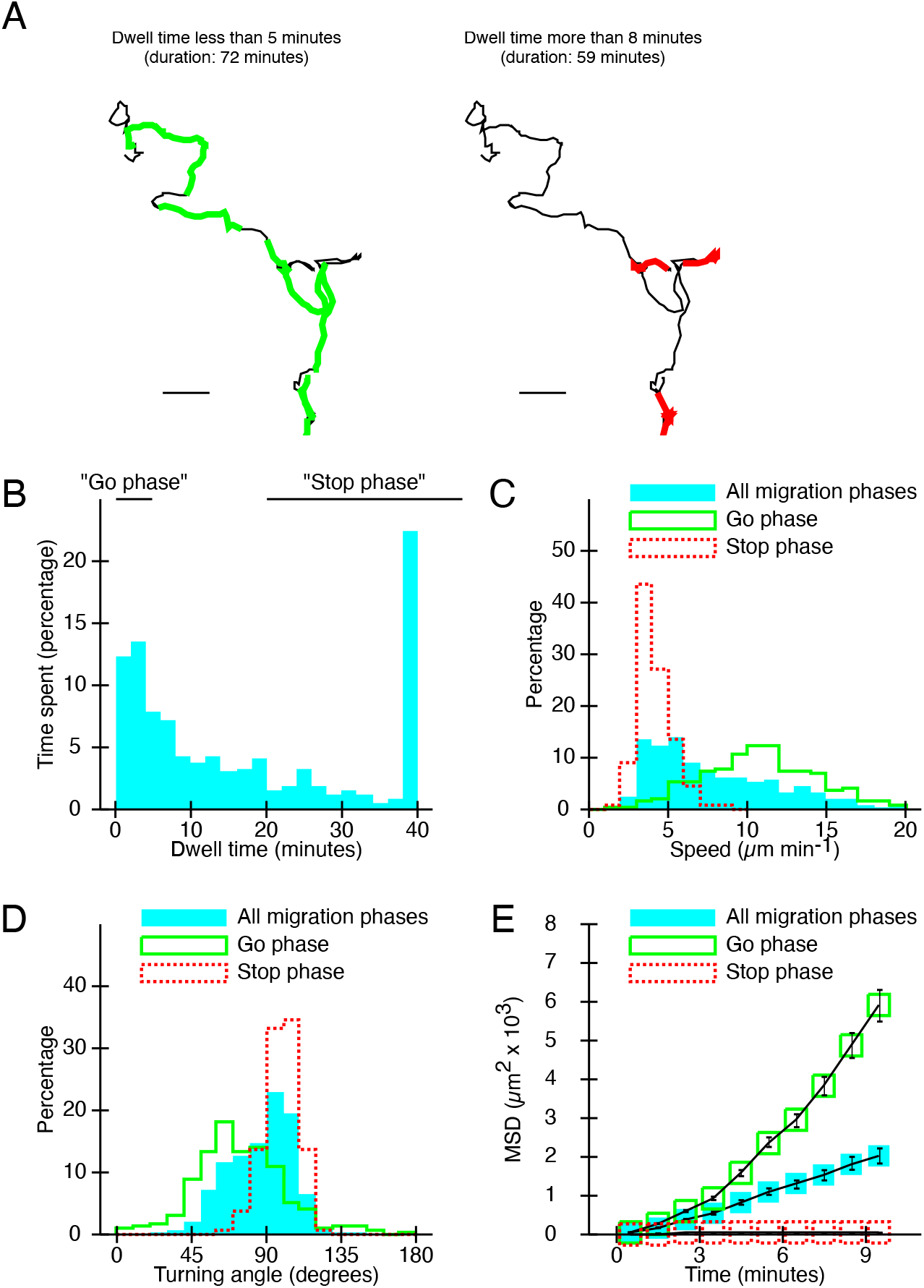
Cytotoxic T cells within influenza-infected lung-parenchyma switch between confined and directional migration modes. C57BL/6 mice received 5 x 10^6^ polyclonal, naïve GFP^+^CD8^+^ T cells obtained from Ubiquitin-GFP mice via the tail vein. The mice were infected one day later with 1 x 10^3^ EID50 X31 influenza. Between day 7 and day 10 post-infection, lungs were isolated and inflated with agarose and transferred into an imaging chamber, followed by live imaging. Generated three-dimensional time-lapse sequences were tracked and analyzed with customized image-analysis algorithms. (A) A representative track, where short dwell times (< 5 minutes, left panel) are highlighted in green, and long dwell times (>8 minutes, right panels) are highlighted in red. (B - E) We analyzed 258 tracks (23,989 frames) obtained from five independent movies, and split these tracks into 1,510 non-overlapping track-segments, each with a displacement of approximately 15 μm. (B) The frequency distribution of dwell times, i.e., the duration of the analyzed track-segments. (C - E) All neighboring 1,092 segments with dwell times < 5 minutes were merged into 290 tracks (4,624 frames; 2,526 minutes) to analyze the “go phase”. All neighboring 125 segments with dwell times > 20 minutes were merged into 110 tracks (14,976 frames; 8,818 minutes) to capture the “stop phase”. The graphs display the frequency distributions of the mean track speeds (C), turning angles (D) and mean squared displacements (E).

Quantification of our entire dataset revealed that most CD8^+^ T cells alternated between either very short (less than five minutes) or long (>= 30 minutes) dwell times (**Figure 5B**). We further merged all neighboring segments with short dwell times to quantify the “go” phase or with long dwell times to quantify the “stop” phase. As expected, T cells moved faster (**Figure 5C**) and straighter (**Figure 5D**) during the go phase than during the stop phase. In addition, the mean squared displacement during the go phase increased super-linearly during observation periods in the range of minutes (**Figure 5E**), indicative of straight ballistic migration. These data show that the dwell time parameter quantifies the switching of CD8^+^ T cells between confined and ballistic migration in the influenza-infected lung.

### High influenza antigen density increases dwell times of lung-infiltrating T cells

We hypothesized that cognate interactions of CD8^+^ T cells with influenza-infected target cells could cause the observed variations of dwell times of CD8^+^ T cells in the influenza-infected lung. To analyze how antigen density affected dwell times, we imaged lungs from X31-infected animals as described previously (Mrass et al., 2017). In addition, we labeled whole tissue with an antibody specific to the H3N2 virus. Visual inspection of CD8^+^ T cells within influenza-high and influenza-low areas of the lung suggested that the dominant migration mode of CD8^+^ T cells in low-influenza lung regions was active translocation (**Figure 6A**, top panel; **Movie 2**). In contrast, in areas with high influenza staining, CD8^+^ T cells appear persistently confined (**Figure 6A**, middle panel; **Movie 3**) or show transitions between confinement and restricted migration (**Figure 6A**, bottom panel; **Movie 4**).

**Figure 6.**
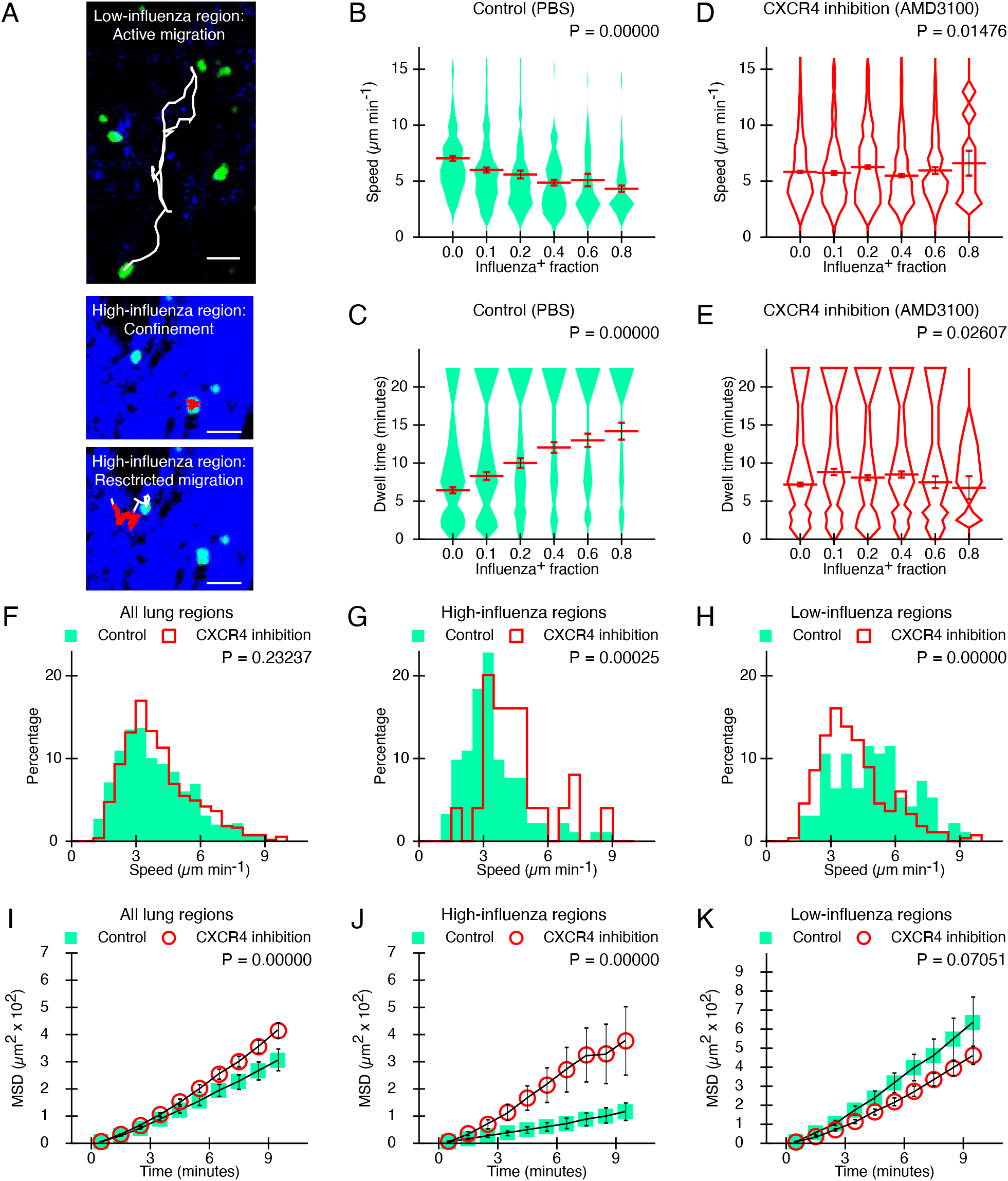
CXCR4 regulates dwell times of lung-infiltrating cytotoxic T cells. (A) C57BL/6 mice received 5 x 10^6^ polyclonal, naïve GFP^+^CD8^+^ T cells obtained from Ubiquitin-GFP mice via the tail vein. The mice were infected one day later with 1 x 10^3^ EID50 X31 influenza. Between day seven and day 10 post-infection, lungs were isolated and inflated with agarose, then stained with a biotinylated antibody specific for H3N2 viral particles and Streptavidin Alexa Fluor 555 secondary. After transfer into an imaging chamber, we captured three-dimensional time-lapse sequences, and tracked migration with customized image-analysis algorithms. Shown are three representative tracks depicting characteristic T cell behavior in regions with high and low influenza densities. White lines show the entire track. Red lines show track segments with dwell times > 20 minutes. Scale bar = 25 μm. (B - K) Mice received naïve GFP^+^CD8^+^ T cells obtained from OT-I x Ubiquitin-GFP mice and were infected with 1 x 10^5^ PFU X31-OVA influenza. From day five post-infection the mice received PBS (control) or the CXCR4 inhibitor AMD3100. Explanted lungs were stained with H3N2 influenza viral particle-specific antibody and AMD3100 was added to the superfused medium at a concentration of 50 μM. After image-acquisition, each pixel of the image volume was designated as influenza antigen-positive or -negative. The fraction of influenza-positive pixels in the image volume surrounding each T cell was quantified by a value between 0 (no positive pixels) and 1 (all pixels are positive). From control lungs, we obtained 414 T cell tracks (29,601 tracked frames) from four independent fields of view, which led to 978 segments, of which 471 were in the go-phase (total duration 1,256 minutes) and 263 were in the stop phase (total duration: 7,891 minutes). From AMD3100-treated lungs, we obtained 706 T cell tracks (52,025 tracked frames) from six independent fields of view, which led to 2,174 segments, of which 1,194 were go segments (total duration 3,233 minutes) and 386 were stop segments (total duration: 11,401 minutes). (B – E) The graphs show mean speeds (B, D) and dwell times (C, E) for segments from control lungs (B, C) or AMD3100-treated lungs (D, E). The lines indicate the mean and the error bars show the standard error of the mean. The bins for the influenza-densities contain 268, 215, 176, 158, 99, 62 values in the control experiment and 810, 366, 544, 359, 86, 9 values in the experiments with AMD3100-treatments. (F – H) Frequency distributions of mean track speeds of all tracks (D, control: 414 tracks, CXCR4 inhibition: 706 tracks), tracks in influenza-dense (E, control: 92 tracks, CXCR4 inhibition: 25 tracks) and influenza-sparse (F, control: 97 tracks, CXCR4 inhibition: 269 tracks) regions. (I – K) Mean squared displacements of all tracks (I), tracks in influenza-dense (J) and influenza-sparse (K) regions. Symbols show the mean of all tracks in each time-bin. Errors show standard error of the mean. Panel J contains 414, 410, 407, 401, 398, 394, 393, 389, 385, 383 values per time bin for control and 706, 700, 695, 693, 692, 688, 684, 680, 671, 668 values for CXCR4 inhibition. Panel K: 92, 92, 92, 90, 90, 90, 90, 89, 89, 89 and 25, 25, 25, 25, 25, 24, 24, 24, 23, 23 values. Panel L: 97, 96, 95, 94, 91, 90, 90, 88, 87, 86 and 269, 267, 265, 264, 264, 263, 261, 261, 257, 257 values. P-values were calculated with the Kruskal-Wallis test (B – E), Mann-Whitney test (F -H) and the Repeated Measures ANOVA test (I – K).

We quantitated the dwell times of cognate OT-I cytotoxic T cells in high-influenza and low-influenza areas in lungs from mice infected with X31-OVA. To systematically quantify the relationship between exposure to cognate antigen and T cell migration behavior, each pixel in the H3N2 channel was classified as influenza-positive or -negative based on the measured intensity level. The fraction of influenza-positive pixels surrounding individual OT-I T cells reflects influenza density. This analysis revealed that OT-I T cells in regions with high influenza density showed reduced speed and increased dwell times compared with OTI T cells in low influenza density areas (**Figures 6B** and **6C**). These results are consistent with findings that T cells can make long-term contact with cognate antigen-presenting target cells *in vivo* (Deguine et al., 2010). An analysis of polyclonal T cells responding to X31 revealed similar results (**data not shown**). These data confirm that high density of influenza antigen increases the dwell time of cognate lung-infiltrating CD8^+^ T cells.

### CXCR4 regulates the stop-signal of T cells within influenza-infected lungs

We then measured CD8^+^ T cell motility patterns in the lungs of influenza-infected mice treated with the CXCR4 inhibitor AMD3100. In contrast to control lungs, where high influenza-antigen density led to confined CD8^+^ T cell migration (**Figure 6B and 6C; Movie 5**), the influenza antigen density had only a marginal effect on OT-I T cell speed and dwell times after CXCR4 inhibition (**Figure 6D** and **6E; Movie 6**). A careful comparison of the motility in regions with different influenza-antigen loads elucidated the basis for the effect of CXCR4 on distinct migration patterns. In areas with high influenza load, CXCR4 inhibition increased the speed (**Figure 6G**) and displacement (**Figure 6J**) of CD8^+^ T cells. In contrast, in low-influenza regions, CXCR4 inhibition reduced the T cells’ speed (**Figure 6H**) and displacement (**Figure 6K**).

Interestingly, when comparing the total CD8^+^ T cell population without regard for antigen density, the effect of CXCR4 on T cell motility was only marginal (**Figures 6F** and **6I**). These results illustrate the importance of performing a separate analysis of CD8^+^ T cell motion depending on antigen density. Quantitation of CXCR4 effects on CD8+ T cell motility in high and low-influenza regions demonstrate that CXCR4 mediates differential effects on CD8^+^ T cell movement within influenza-infected lungs depending on antigen density. In antigen-dense areas, CXCR4 promotes CD8^+^ T cells arresting on target cells, while CXCR4 enhances the motility of CD8^+^ T cells in areas with low antigen.

### Inhibition of CXCR4 improves recovery of influenza-infected mice

We then determined how CXCR4 inhibition in the effector phase of the T cell response influenced the outcome of influenza infections. Treatment of X31-infected mice with AMD3100 from day five post-infection changed the kinetics of recovery of lost weight (**Figure 7A**). During the early recovery phase between days seven and eight, inhibition of CXCR4 slightly delayed recovery of lost weight (**Figure 7A** and **data not shown**). Inhibition of CXCR4 also delayed clearance of replication-competent virus from lungs on day seven (**Figure 7B**), suggesting ineffective killing of infected epithelial cells by cytotoxic T cells.

**Figure 7.**
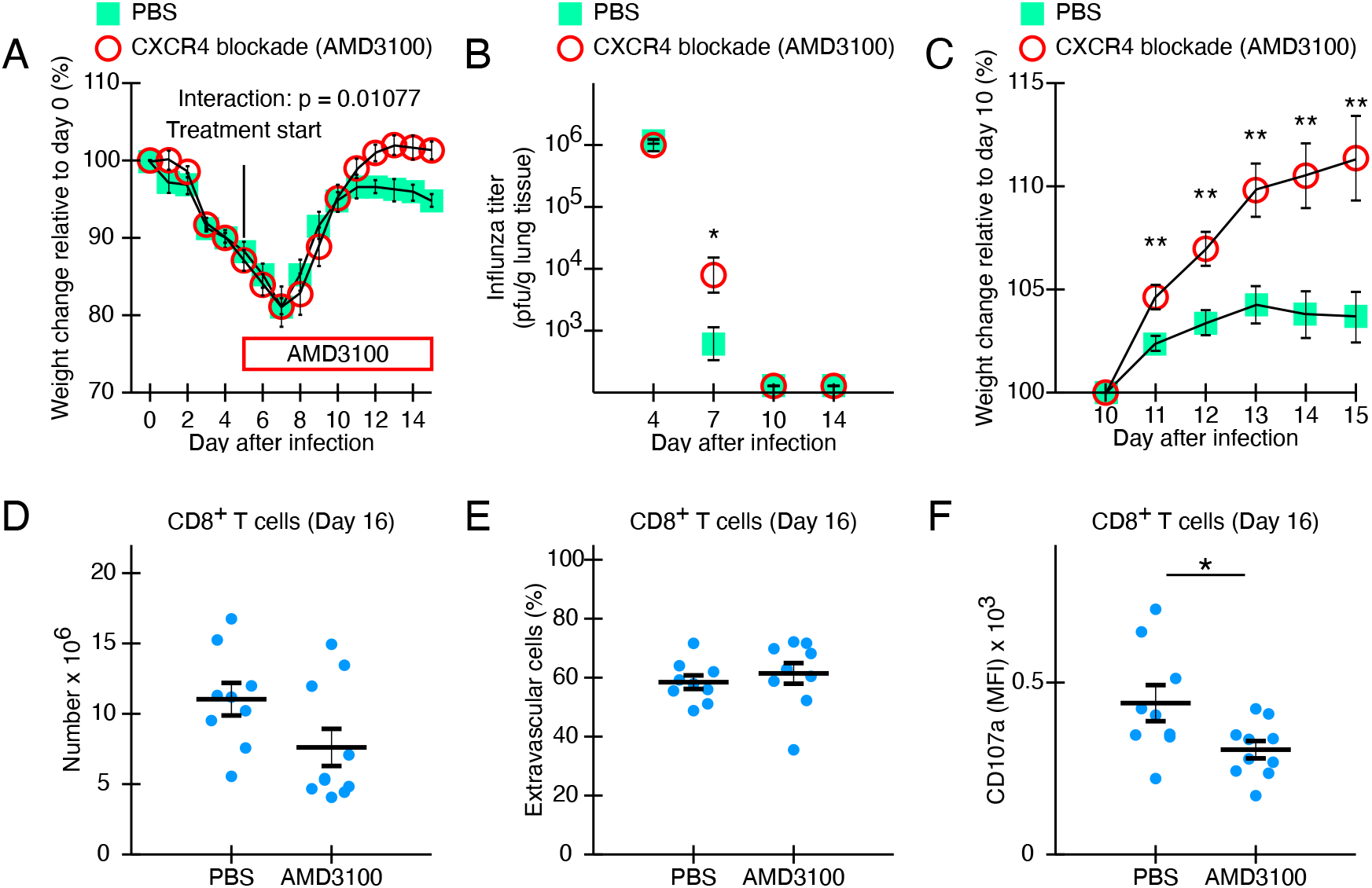
Inhibition of CXCR4 promotes recovery of influenza-infected mice. We infected mice with 1 x 10^5^ PFU X31-OVA influenza and measured weights until day 15 after infection. Starting on day five after infection, mice received PBS (control) or the CXCR4-inhibitor AMD3100 by intraperitoneal injection every 12 hours. (A) Weight percentage relative to start obtained from a representative experiment with five mice in each group. Lines show means and error bars indicate standard errors of the mean. The p-value for the interaction was calculated with the two-way ANOVA. (B) Mean influenza titer obtained from lungs of infected mice (four mice per group at each time point). Error-bars show the standard error of the mean. The p-value was calculated with the Student’s t-test. (*; P-value < 0.05). (C) Weight changes relative to day 10 for each mouse from pooled data from three independent experiments. Each group contains measurements from 17 mice. P-values for each timepoint were calculated with the Student’s t-test (**: P-value < 0.005). (D- F) Single-cell suspensions were stained to detect extravascular CD8^+^ T cells and measure the surface level of CD107a. Symbols show individual mice. The graphs show data pooled from two independent experiments.

When we assessed mouse weight after viral clearance between days 10 and 15 post-infection, we found that AMD3100-treated mice gained more weight than control mice (**Figure 7C**). To determine whether changes in CD8^+^ T cell function correlated with the increased weight gain after viral clearance, we assessed the effect of CXCR4 on effector CD8^+^ T cells on day 16 post-infection. Flow cytometry analysis revealed similar numbers (**Figure 7D**) and extravasation (**Figure 7E**) of CD8^+^ T cells in the lung tissue in AMD3100-treated and untreated animals. In contrast, inhibition of CXCR4 reduced degranulation measured by CD107a in lung-infiltrating CD8^+^ T cells (**Figure 7F**) at day 16 post-infection.

These data suggest that CXCR4 affects multiple aspects of immune responses throughout influenza infection. CXCR4 enhances virus clearance on day seven post-infection, when the CD8^+^ T cell response is at its peak. However, after viral clearance, CXCR4 increases continued T cell degranulation, and animals remain at lower weights during recovery from influenza infection. These differential effects of CXCR4 throughout the immune response to influenza suggest that CXCR4 plays a role in balancing protective immunity and immune-mediated tissue damage during viral infection.

## Discussion

During respiratory tract infections, cytotoxic T cells boost protective immunity by eliminating infected cells but can also exacerbate disease severity by increasing immunopathology (Duan and Thomas, 2016; Hufford et al., 2015; Rutigliano et al., 2014). However, our understanding of the mechanisms that regulate cytolytic T cell function within the lung remains incomplete. In this study, we used a murine model of influenza lung infection to investigate the effect of CXCR4 on the functional activity of cytotoxic T cells in the lung after infection. Our principal finding is that CXCR4 on effector CD8^+^ T cells promotes T cell activation, degranulation, and interaction with target cells in the influenza-infected lung.

In addition, we identify opposing roles for CXCR4 depending on influenza antigen density. In high-antigen areas, CXCR4 is essential for triggering the “stop signal” of CD8^+^ effector T cells, which enhances T cell activation. In contrast, in regions with low levels of antigen, CXCR4 enhances migration. This “seek and destroy” function of CXCR4 works by increasing migration in antigen-sparse regions and improving interaction in antigen-dense areas. This combined effect of CXCR4 will likely enhance the total number and duration of interactions between cytotoxic T cells and antigen-presenting target cells in the lung environment.

Chemokines are a group of molecules best known to regulate the entry of immune cells from the vasculature into lymphoid and peripheral tissues, including influenza-infected lungs (Wareing et al., 2004). Chemokines also regulate T cell motility *in situ* within tissues to promote the enrichment of cytotoxic T cells in specific tissue subregions, such as infectious foci within the skin and the epithelial surface of the influenza-infected trachea (Ariotti et al., 2015; Germain et al., 2012; Lambert Emo et al., 2016; Lim et al., 2015; Overstreet et al., 2013; Prizant et al., 2021). These reports showed that modest effects on T cell migration parameters such as speed can cause significant changes in intra-tissue positioning and modulate T cell function (Ariotti et al., 2015; Boyd et al., 2020). Our findings also align with previous reports showing increased T cell migration in the influenza-infected lung after viral clearance. In addition, they suggest that increases in the motility of T cells in lung regions with low influenza density can help T cells to reach virally infected target cells within influenza antigen-dense areas (Lambert Emo et al., 2016; Matheu et al., 2013).

We also identify a new role for CXCR4 in regulating CD8^+^ effector T cell interactions within the lung. First, we show that CXCR4 promotes calcium flux, the release of cytotoxic granules, and interaction with antigen-bearing target cells in an *in vitro* system using CXCR4 knockout T cells, where T cells initiate interactions independent of their migratory activity. Thus, our results show that CXCR4 has T cell-intrinsic effects on CD8^+^ T cell activation and effector function. Moreover, our data show that CXCR4 enhances T cell dwell times within the lung and increases the magnitude of degranulation in individual T cells. This evidence suggests that CXCR4 has a similar stimulatory role on T cell interactions *in vivo*. Our findings add insight to previous studies that have shown recruitment of CXCR4 into the immunological synapse *in vitro* (Bromley et al., 2000; Molon et al., 2005; Nanki and Lipsky, 2000; Peacock and Jirik, 1999). Together, our results indicate that CXCR4 strengthens cytotoxic T cell interactions both *in vitro* and *in vivo*.

Our studies took advantage of a well-established inhibitor of CXCR4, AMD3100. This drug is suitable for treating human patients and animals and effectively inhibits CXCR4 in the influenza-infected trachea and other tissues (Collins et al., 2019; Feig et al., 2013; Molon et al., 2005; Uy et al., 2012). This approach enabled us to target CXCR4 primarily during the effector phase but not at the priming phase of the CD8^+^ T cell response. We found that inhibition of CXCR4 after the T cell priming phase showed little to no effect on T cell migration into the influenza-infected lung. In contrast, CXCR4 inhibition slightly increased weight loss between days seven and eight, i.e., when CD8^+^ effector response is maximal. Inhibition of CXCR4 also increased weight gain during the late recovery phase. We also found that CXCR4 inhibition led to decreased CD8^+^ effector T cell function at late time points, showing a role for CXCR4 in the downregulation of CD8^+^ effector responses even after viral clearance.

A comparison of CXCR4 to the immune-checkpoint molecule PD-1 reveals several contrasts. First, PD-1 knockout mice lose less weight during early viral lung infections than wild-type animals but recover more slowly after clearance of the virus (Erickson et al., 2012). This finding is the opposite of what we observe after the inhibition of CXCR4. Second, PD-1 antagonizes the stop-signal of T cells that migrate in the pancreas (Fife et al., 2009), while CXCR4 strengthens the stop signal. These findings suggest that CXCR4 and PD-1 may antagonistically control the strength of the stop-signal of migrating T cells in tissues, leading to opposite biological outcomes.

In conclusion, the results of the present study show that CXCR4 is crucial for the activation, degranulation, and migration of cytotoxic T cells in influenza-infected lungs. Other chemokine receptors have distinct effects on T cell function. For example, deficiency of the chemokine receptor CXCR3 delayed recruitment of lymphocytes into influenza-infected lungs, which was associated with delayed clearance of the virus (Gilchuk et al., 2016; Kohlmeier et al., 2009; Wareing et al., 2004). These findings extend our understanding of how the chemokine receptor CXCR4 regulates multiple aspects of effector CD8^+^ T cell function. In addition to regulating migration to and within tissues, CXCR4 can also strengthen the interaction, activation, and degranulation of cytotoxic T cells within infected peripheral tissue. Post viral clearance, inhibiting CXCR4 might mitigate CD8^+^ T cell-mediated immunopathology after the acute viral phase of infection.

## Supporting information

Mrass Supplemental Movie 1

Mrass Supplemental Movie 2

Mrass Supplemental Movie 3

Mrass Supplemental Movie 4

Mrass Supplemental Movie 5

Mrass Supplemental Movie 6

## Movie Legends

**Movie 1. Lung-infiltrating cytotoxic T cells show changes in dwell time over time.**

CD8^+^ T cells (GFP^+^CD8^+^ from B6Ub-GFP animals) were imaged in the lungs of mice infected with X-31 influenza A virus between days seven to 10 post-infection. The pink square marks the T cell’s centroid over time. The white line indicates the T cell tracking. The green lines highlight migration phases with less than five minutes dwell time, and the red lines highlight migration phases with more than eight minutes. The lower left corner shows the time after image capture initiation, and the lower right corner shows the scale bar.

**Movie 2. A cytotoxic T cell in a lung region with a low level of influenza-antigen shows active migration.**

CD8^+^ T cell imaging and annotation are the same as in Movie 1 with the addition of influenza-antigen staining (blue) as described in Figure 6 and Materials and Methods.

**Movie 3. A cytotoxic T cell in a lung region with a high level of influenza antigen is confined.**

CD8^+^ T cell imaging and annotation are the same as in Movies 1 and 2, except that the red line highlights migration phases with a dwell time above 20 minutes.

**Movie 4. A cytotoxic T cell in a lung region with a high level of influenza antigen shows restricted migration.**

CD8^+^ T cell imaging and annotation are the same as in Movie 3.

**Movie 5. Cytotoxic T cells in a lung region with high levels of influenza antigen show low migratory activity.**

CD8^+^ T cells were imaged in lungs of X-31 infected mice as described in Figure 6. CD8^+^ T cells (green) move through a lung region with a high level of influenza antigen (blue). The white lines represent T cell tracks, and the blue crosses indicate centroid positions. The left lower corner shows the time after image capture initiation, and the right lower corner shows the scale bar.

**Movie 6. After inhibition of CXCR4, cytotoxic T cells in a lung region with high levels of influenza antigen show increased migratory activity.**

CD8^+^ T cell from lungs of animals treated with AMD3100 from day five post-infection were imaged. Explanted lungs were also superfused with medium containing AMD3100 during imaging (Materials and Methods). Annotation is the same as in Movie 5.

## Acknowledgements

This work was supported by funding from the following: NIH 1R01AI097202 (JLC), the Spatiotemporal Modeling Center (P50 GM085273), the Center for Evolution and Theoretical Immunology 5P20GM103452 (JLC), AIM CoBRE at UNM HSC NIH P20GM121176 (JLC; PM), NIH 5 T32 AI007538-19 (JRB), an Institutional Development Award (IDeA) from NIGMS P20GM103451 (DJT) and RL5GM118969 (DJT); institutional support from dedicated health research funds from UNM SOM (JLC), Women in Science Award from UNM, DARPA/AFRL FA8650-18-C-6898 (JLC) and in part institutional support from Dedicated Health Research Funds from the UNM School of Medicine (JLC and PM); support by the University of New Mexico Comprehensive Cancer Center Support Grant NCI P30CA118100 (PM and IK). I.K. was supported by a Liz Tilberis Early Career Development Award from the Ovarian Cancer Research Alliance, and additional support was provided by the and the Flow cytometry and Microscopy Core at UNMCCC.

## Methods

### Mice and reagents

C57BL/6 (Jackson Laboratories) mice or CD4-Cre:CXCR4 mice were used. CD4-Cre (Jackson Tg(Cd4-cre)1Cwi/BfluJ) were bred to CXCR4^Flox/Flox^ mice (Jackson Laboratories B6.129P2-Cxcr4^tm2Yzo^/J) to generate mice with T cells deficient in CXCR4 (CXCR4^KO^). As controls, we used littermates lacking the CD4-Cre transgene (CXCR4^WT^). We confirmed CXCR4-deficiency by genetic screen and flow cytometry. All mice were maintained in a specific pathogen-free environment in barrier facilities at the University of New Mexico School of Medicine in Albuquerque, NM, and conform to the principles outlined by the Animal Welfare Act and the National Institutes of Health guidelines and approved by the IACUC animal use committee. Males and females were used at between 8-20 weeks. All experimental protocols were approved by the IACUC animal use committee at UNM Health Sciences Center in accordance with relevant guidelines and regulations (IACUC protocol #s: 18-200797-B-HSC; 16-200497-HSC; 19-200892-HSC). All animal work was performed and reported according to ARRIVE guidelines.

### Reagents including antibodies

We used the following reagents: CXCR4 inhibitor AMD3100 (commercially called Plerixafor) (Tocris, Cat#: 3299), Cal-520 (AAT Bioquest, Cat#: 21130), Streptavidin Alexa Fluor 555 (ThermoFisher, Cat#: S32355). We used the following antibodies: H3N2 virion antibody (ViroStat, Cat#: 1317); anti-mouse CD107A, Alexa Fluor488 (Clone 1D4B; BioLegend, Cat#: 121608); anti-mouse APC/Cyanine7 mouse CD107a Antibody (Clone 1D4B; BioLegend, Cat#: 121616); anti-CD184 (CXCR4) Monoclonal Antibody (2B11), PE (Invitrogen, Cat#: 12-9991-82); PE anti-mouse CD183 (CXCR3) Antibody (R&D, Cat# 126505); anti-Mouse CD3, PE (Clone: 17A2; BD Pharmingen; Cat#: 555275); anti-Mouse CD3e, PerCP-Cy5.5 (Clone: 145-2C11, eBioscience, Cat#: 45-0031-82); anti-Mouse CD3, Alexa Fluor 647 (Clone: 17A2; BD Pharmingen, Cat#: 557869); anti-Mouse CD8a, APC-eFluor780 (Clone: 53-6.7; eBioscience, Cat#: 47-0081-82); anti-Mouse CD4, PE-Cyanine7 (Clone: GK1.5; eBioscience, Cat#: 25-0041-82).

### Influenza infection of mice

We infected mice intranasally with 30 μl of PBS containing either 1 x 10^3^ EID50 of the Influenza A X31 strain (Charles River) or 1 x 10^5^ PFU of Influenza A X31-OVA (kindly provided by Dr. Paul Thomas). X31 contains internal genes for A/Puerto Rico/8/1934 (H1N1) with HA and NA from A/Aichi/2/1968 (H3N2). X31-OVA additionally encodes a peptide from OVA (SIINFEKL) in the NA stalk. During infection, mice were under anesthesia induced with 90 mg/kg ketamine and 8.1mg/kg xylazine. This approach ensures sedimentation of the virus into the lower respiratory tract.

### Measurement of influenza titer from infected lungs

Virus titers were determined by plaque assay on MDCK cells as described (dx.doi.org/10.17504/protocols.io.n2bvj63bxlk5/v1). Briefly, we harvested lungs from infected mice, and stored them in −80 degrees after snap-freezing in liquid nitrogen. Whole lung homogenates were prepared in virus growth media (VGM; DMEM, 23 mM HEPES, 0.2% BSA, 1X penicillin/streptomycin, 2 μg/ml TPCK-trypsin) using a Tissue Lyser II (Qiagen) with the following program: 3 cycles at 30 Hz for 30 s. The concentration of virus in the homogenate was determined by infecting MDCK cells at 37°C, 5% CO2 in VGM plus 1.2% avicel overlay. Plaques were visualized 4 days after infection by crystal violet staining, and titer calculated per g of lung tissue. Lungs without recoverable virus were set to the limit of detection.

### Adoptive transfer of CD8^+^ T cells into recipient mice

Mice received 1 x 10^6^ polyclonal naïve GFP^+^CD8^+^ T cells from Ubiquitin-GFP mice or 1 x 10^4^ GFP^+^OT-I T cells from OT-I x Ubiquitin-GFP mice before infection with influenza X31. Single-cell suspensions obtained from spleens were processed with the CD8a^+^ T Cell Isolation Kit (Miltenyi Biotec; Cat#: 130-104-075) and transferred via the tail vein into recipient mice.

### Analysis of lung samples with flow cytometry

Five minutes before tissue harvest, mice received 3 μg of an anti-CD3 antibody (clone 17A2) via the tail vein. We obtained lung single cell suspensions with the Mouse Lung Dissociation Kit (Miltenyi Biotec). We stained the cells with an antibody solution to detect CD8^+^ T cells, CXCR3, CXCR4 and CD107a, fixed them with a 2% paraformaldehyde solution, and quantified them with the LSR Fortessa. Exported fcs files were analyzed with FlowJo (Becton Dickinson).

### Measurement of calcium mobilization

We generated effector CD8^+^ T cells from CD4-Cre : CXCR4^Flox/Flox^ mice where the CXCR4 gene is knocked-out in all T cells (CXCR4^KO^). As controls, we used littermates that did not express CD4-Cre (CXCR4^WT^). We isolated CD8^+^ T cells from spleens with the CD8a^+^ T Cell Isolation Kit (Miltenyi Biotec) and plated them on dishes coated with an antibodies for CD3 (2 μg/ml; clone 145-2C-11; BioXCell) and CD28 (1 μg/ml; clone PV-1; BioXCell). We cultured the T cells in DMEM medium (Gibco, #11995-065) or RPMI (Gibco, Cat#: 22400-105) supplemented with 0.6 mM beta-mercaptoethanol, 5% fetal calf serum, 10 mM HEPES (Gibco; Cat#: 15630-080), MEM non-essential amino acids (Gibco; Cat#: 11140-0500) and Pen/Strep (Gibco; Cat#: 15140-122), and every other day added fresh media supplemented with human interleukin-2 (20 units/ml; TECINTM, Teceleukin). We used the cells four to seven days after stimulation. To quantify calcium mobilization, we loaded cytotoxic T cells for 30 minutes with 4 μM Cal-520 (AAT Bioquest), and transferred 2 x 10^4^ cells per well into flat-bottom 96 well plates coated with a CD3 antibody. We immediately transferred loaded 96 well plates into the Incucyte Imaging System (Sartorius). We generated time-lapse sequences with a frame-rate of one image per minute, while the T cells were sedimenting. We quantified mean fluorescence of Cal-520 for single T cells starting from contact initiation with the plastic dish.

### Measurement of T cell degranulation *in vitro*

We prepared cytotoxic T cells like for the calcium mobilization assay. On the day of the experiment, we added 2 x 10^4^ of the expanded cytotoxic T cells into 96 wells coated with CD3-antibody. Some wells contained 50 μM of the CXCR4 inhibitor AMD3100 (Tocris). We centrifuged the 96 well plates for 1 minute with 1,350 rpm to sediment the cells to the bottom of the well. After two hours of incubation, we stained the T cells with antibodies for CD107a (Clone 1D4B; BioLegend) or CXCR4 (Clone 2B11; Invitrogen), and quantified their fluorescence level with the LSR Fortessa. Percentages of CD107a+ T cells were calculated with FlowJo.

### Measurement of T cell interactions *in vitro*

We generated OVA-specific cytotoxic T cells by pulsing splenocytes from Ubiquitin-GFP x OT-I mice with 1 μg ml^-1^ SIINFEKL peptide. We then expanded the T cells as described for the calcium mobilization experiment. One day before the experiment, we added 3 x 10^3^ ID8-OVA-RFP cells into 96 wells so that they formed a subconfluent target cell layer. On the day of the experiment, we added 2 x 10^4^ cytotoxic OT-I T cells into the 96 wells. After several hours, we gently added a prewarmed paraformaldehyde solution into each well to achieve a final concentration of 2%. We continued incubation for another 15 minutes to fix interactions between the cytotoxic T cells and the adherent ID8-OVA cells. After washing and storing the fixed cells in PBS, we captured the GFP^+^ OT-I T cells and RFP^+^ ID8-OVA cells with a Zeiss LSM900 microscope. We segmented individual ID8-OVA cells and T cells and then quantified how many T cells interacted with single ID8-OVA cells.

### Treatment of mice with the CXCR4 inhibitor AMD3100

Starting from day five after infection with influenza, we injected 200 μg of AMD3100 in 200 μl PBS by intraperitoneal infection, two times a day until sacrifice.

### Live tissue imaging of influenza-infected lungs

After euthanizing the mice, we inflated the lungs by injecting 1 ml of a 37 degree 2% low melting agarose solution via a catheter inserted in the trachea (Sigma-Aldrich, Cat#: A0701). We solidified the agarose by pouring a cold PBS solution (4 degrees Celsius) over the inflated lungs. For co-visualization of T cells and influenza-infected lung regions, we incubated the lungs for 30 minutes with a biotinylated anti-H3N2 influenza viral particle antibody in T cell culture medium (ViroStat, Cat#: 1317), followed by washing and staining with Streptavidin Alexa Fluor 555 (ThermoFisher; Cat#: S32355). We then transferred the lungs in a biosafety cabinet into a POC-R imaging chamber (LaCon) and imaged the lungs with a Zeiss LSM800 Airyscan Confocal Microscope. Due to the transparency of the prepared lung tissue, it was possible to visualize at tissue depths of up to 60 μm. We continuously perfused the chamber with media bubbled with a 95% oxygen 5% CO2 gas mixture and maintained the temperature of the media at 37 degrees. In some experiments, we used a Prairie Ultima Two-photon microscope or a Zeiss LSM 510 microscope, which captured equivalent T cell behavior.

### T cell tracking

We quantified T cell migration and interaction with ImageTracker, a collection of MatLab classes (https://github.com/pmrass/ImageTracker). First, we generated T cell tracks by computational algorithms, followed by manual refinement that captured the three-dimensional position of T cells every thirty seconds. The mean track speed is the sum of the magnitude of all vectors between consecutive time frames divided by the total track duration. Next, we calculated turning angles between two successive track vectors ranging from 0 to 180 degrees. Finally, for the mean squared displacement of individual cells, we defined time intervals of interest, then calculated for each interval a single value by averaging the displacements from all appropriate time intervals of the cell track.

### Algorithm to recognize confined “stop” and active “go” T cell migration phases

We split parent tracks into a series of track segments to detect T cells during “stop” and “go” phases. We started from the beginning of each track and split off the first segment when the displacement from the origin exceeded 15 μm and repeated this procedure until the end of each track. Thus, we converted each parent track into a series of non-overlapping track segments, each with a displacement of approximately 15 μm. We called the duration of each track “dwell time” because it represents the time that cells dwell in a confined tissue region, without spatial relocation >15 μm. The duration of the dwell times determined the T cells’ migration classification.

### Measurement of T cell migration as a function of influenza-density

We classified each pixel within the generated image volumes as influenza-positive or - negative, based on the thresholding of influenza-stained images. Then, we quantified the percentage of influenza-positive pixels surrounding a T cell at each time frame as a measure of influenza-antigen density.

